# Volatile anesthetics inhibit presynaptic cGMP signaling to depress presynaptic excitability in rat hippocampal neurons

**DOI:** 10.1101/2022.01.03.474845

**Authors:** Iris Speigel, Vanessa Osman, Hugh C Hemmings

## Abstract

Volatile anesthetics alter presynaptic function including effects on Ca^2+^ influx and neurotransmitter release. These actions are proposed to play important roles in their pleiotropic neurophysiological effects including unconsciousness and amnesia. The nitric oxide and cyclic guanosine monophosphate (NO/cGMP) signaling pathway has been implicated in presynaptic mechanisms, and disruption of NO/cGMP signaling has been shown to alter sensitivity to volatile anesthetics *in vivo*. We investigated NO/cGMP signaling in relation to volatile anesthetic actions in cultured rat hippocampal neurons using pharmacological tools and genetically encoded biosensors of cGMP levels. Using the fluorescent biosensor cGull we found that electrical stmulation-evoked NMDA-type glutamate receptor-independent presynaptic cGMP transients were inhibited −33.2% by isoflurane (0.51 mM) and −23.8% by sevoflurane (0.57 mM) (p<0.0001) compared to a stimulation without anesthetic. Isoflurane and sevoflurane inhibition of stimulation-evoked increases in presynaptic Ca^2+^ concentration, measured with synaptophysin-GCaMP6f, and synaptic vesicle exocytosis, measured with synaptophysin-pHlourin, were reduced by in neurons expressing the cGMP scavenger sponGee. This reduction in anesthetic effect was recapitulated by inhibiting HCN channels, a cGMP-modulated effector that can facilitate glutamate release. We propose that volatile anesthetics depress presynaptic cGMP signaling and downstream effectors like HCN channels that are essential to presynaptic function and excitability. These findings identify a novel mechanism by which volatile anesthetics depress synaptic transmission via second messenger signaling involving the NO/cGMP pathway.

## Introduction

Nitric oxide (NO) is a gaseous messenger that is generated from L-arginine by nitric oxide synthase (NOS) in a calcium (Ca^2+^)-dependent manner. The NO signaling system is widely expressed and pleiotropic in function, with prominent roles in the nervous system that include several forms of synaptic modulation. The primary downstream effector of NO is soluble guanylyl cyclase (sGC), which converts GTP into the second messenger cyclic GMP (cGMP), which itself acts on multiple downstream targets including ion channels and signaling proteins ^1^. Several lines of evidence suggest a relationship between NO signaling and potency of volatile anesthetics^2^. However, the precise mechanisms have not been identified in part due to the complexity of NO/cGMP signal transduction pathways, with conflicting results reported in previous studies^3, 4^

Pharmacological studies have shown increased sensitivity to loss of righting reflex with inhibition of NOS and sGC, which suggests a contribution of reduced NO signaling to anesthetic-induced hypnosis^4–6^. However, knockout of the sGC-1α isoform in mice was recently shown to reduce sensitivity to sevoflurane anesthesia as measured by both loss and return of righting reflex, suggesting that sGC1α-derived cGMP contributes to volatile anesthetic actions^7^. Although one study found that knockout of neuronal NOS (nNOS) did not alter sensitivity to isoflurane anesthesia^4^, nNOS inhibitors can increase sensitivity to volatile anesthetic-induced loss of righting reflex, suggesting nonspecific pharmacological and/or genetic effects. A number of studies have reported volatile anesthetic effects on brain NO/cGMP concentrations. Halothane, isoflurane, and sevoflurane all increase cerebral NO^8–10^ and cGMP^7, 11, 12^ concentrations, although decreased NO and cGMP levels have also been reported^13^.

The multiple behavioral outcomes resulting from manipulating nNOS signaling may be related to off-target effects of volatile anesthetics (i.e. on the cardiovascular system) that complicate interpretation of global gene knockout models. Thus it is unclear whether changes in neuronal NO/cGMP signaling are causal or incidental to anesthetic effects in these animal models. Studies *in vitro* have also yielded conflicting findings. For example, in cultured rat cortical neurons, isoflurane has been reported to inhibit^14, 15^ or enhance^14, 15^ glutamate agonist-induced increases in cGMP. Taken together, the relationship between volatile anesthetic actions and cGMP production remains unclear, both *in vivo* and *in vitro*.

A technical barrier to elucidating NO/cGMP signaling pathways is that intracellular cGMP dynamics have been difficult to measure. Studies have implicated cGMP as a regulator of neurotransmitter release via downstream effectors^1^, however none have measured changes in presynaptic cGMP levels directly. Previous studies of cGMP signaling have relied on biochemical measurements of net changes in cell lysates, so anesthetic actions on presynaptic cGMP in real time are unknown. Given that volatile anesthetics inhibit neurotransmitter release via several mechanisms, we hypothesized that alterations in cGMP may contribute to volatile anesthetic actions on presynaptic function. To investigate the potential role of NO/cGMP signaling in volatile anesthetic effects on neurotransmitter release, we took advantage of recently developed genetically encoded approaches to measure and manipulate their presynaptic actions, including the cGMP biosensor green cGull^16^ and the cGMP scavenger sponGee^17^.

## Methods

All experimental procedures were approved by the Weill Cornell Medical College Institutional Animal Care and Use Committee and conformed to National Institutes of Health (Bethesda, MD, USA) Guidelines for the Care and Use of Animals and adhered to ARRIVE guidelines where applicable.

### Primary neuron culture and transfection

Hippocampi were dissected from postnatal rats (0-1 days old, both sexes). Cells were dissociated and plated onto coverslips as described^18^. Transfection was performed on day 6 or 7 *in vitro* (DIV6-7) using Ca^2+^ phosphate-mediated gene transfer. Live-cell imaging was performed between DIV16-20. For each experiment, neurons were derived from at least three separate culture preparations.

### Plasmids

Green cGull was a gift from Tetsuya Kitaguchi (Tokyo Institute of Technology, Kanagawa, Japan; Addgene plasmid # 86867). SponGee was a gift from Xavier Nicol (French National Centre for Scientific Research, Paris, France; pCX-SponGee-mRFP, Addgene plasmid # 134775). Synaptophysin-pHluorin (syn-pH) was a gift from Stephen Heinemann and Yongling Zhu (Northwestern University, Evanston, IL; pcDNA3-SypHluorin 2x, Addgene plasmid # 37004). VAMP-mCherry and synaptophysin-GCaMP6f were gifts from Timothy Ryan (Weill Cornell Medicine, New York, NY)^19^.

### Live-cell Imaging

Experiments were performed on a ZEISS Axio Observer Z1 widefield fluorescence microscope with filter cubes and LEDs for eGFP and RFP detection (Colibri 7 LED illumination modules 475nm and 555nm, with 38HE and 90HE filter cubes, all Zeiss, Oberkochen, Germany) using an Andor iXon1 EMCCD camera (Belfast, Ireland). Synaptic vesicle (SV) exocytosis signals were acquired at 10 Hz, Ca^2+^ signals at 20 Hz, and cGMP at 40 Hz. Coverslips were mounted in a closed field stimulation perfusion chamber with a total volume of 263 μl. Solutions were maintained at 37.0 ± 0.2°C by an in-line heater and imaging chamber heater, and were perfused at 1 ml/min using a custom system and a multibarrel manifold (Warner Instruments, Hamden, CT, USA). The standard buffer was Tyrode’s solution (119 mM NaCl, 2.5 mM KCl, 2 mM CaCl2, 2 mM MgCl2, 25 mM HEPES buffered to pH 7.4, 30 mM glucose) containing 10 μM 6-cyano-7-nitroquinoxaline-2,3-dione (CNQX) and 50 μM D,L-2-amino-5-phosphonovaleric acid (AP5) (both from Tocris, Bristol, UK) to block recurrent glutamatergic signaling.

Neurons expressing cGull were identified by resting green fluorescence, and presynaptic boutons were identified by co-transfected VAMP-mCherry red fluorescence. Neurons coexpressing syn-pH or syn-GCaMP6f were identified by co-transfected mCherry- or mRFP-conjugated sponGee.

Action potentials (APs) were stimulated by trains of 20 stimuli generated by an electrical field pulse generator (Master-9, A.M.P.I., Jersalem, Israel) at 20 Hz, with a stimulus isolator (Model A385, World Precision Instruments, Sarasota, FL, USA) and platinum/iridium bath electrodes built into the imaging chamber yielding a field of 10 V/cm^2^. A maximum of one imaging experiment was acquired per coverslip.

### Volatile anesthetic delivery

An experimental solution of ~0.51 mM isoflurane dissolved in Tyrode’s buffer was prepared daily from a saturated stock solution of 12 mM; this corresponds to ~1.4 MAC (minimum alveolar concentration) in mice^20^. An experimental solution of ~0.0.57 mM sevoflurane was prepared from a saturated stock solution of 6 mM; this corresponds to ~1.2 MAC^20^. Isoflurane or sevoflurane solutions were focally perfused from gas-tight glass syringes into the imaging chamber for 5 min before imaging to allow equilibration. At the conclusion of each experiment, a perfusate sample was taken from the chamber for determination of delivered concentration by gas chromatography (Shimadzu GC-2010 Plus, Kyoto, Japan) with external standard calibration. The reported values of 0.51 ± 0.09 mM isoflurane and 0.57 ±0.13 mM sevoflurane reflect averaged measurements from all bath samples collected.

### Drugs

Isobutylmethylxanthine (IBMX, 100 μM) was purchased from Thermo Scientific (PHZ1124, Waltham, MA, USA). N-ω-nitro-L-arginine (L-NNA, 100 μM), and ivabradine hydrochloride (30 μM) were from Sigma-Aldrich (B1381, N5501, and SML0281, St. Louis, MO, USA). Spermine-NONOate (50 μM) was from Cayman Chemical (82150, Anne Arbor, MI, USA). N5-(1-Imino-3-butenyl)-L-ornithine (L-VNIO, 100 nM) was from Enzo Life Science (ALX-270-216-M005, Farmington, NY, USA). KT-5823 (500 nm) was from Tocris Bioscience (1289, Bristol, UK). IBMX, ivabradine and KT-5823 were perfused for 15 min prior to recordings. L-VNIO and L-NNA were perfused for 20 min prior to recordings^21^.

### Image and statistical analysis

Live-cell imaging recordings were analyzed in ImageJ (rsb.info.nih.gov/ij) using the plugin TimeSeries Analyzer (rsb.info.nih.gov/ij/plugins/time-series.html) to measure fluorescence over time. For presynaptic Ca^2+^ measurements, fluorescence was analyzed within 2 μm diameter regions of interest (ROIs), each containing a visually identifiable synaptic bouton. ROIs were selected based on their peak response and latency to control stimulation in Tyrode’s buffer. Time-series data were exported into MATLAB for further analysis with custom scripts. For each ROI, fluorescence was calculated as ΔF/F_0_ = ((F-F_0_)/ F_0_). Baseline fluorescence (F_0_) was averaged from the 10 frames immediately prior to stimulus onset, and peak fluorescence (F) was averaged from the three consecutive frames with the highest values following stimulation. ROIs with signal-to-noise ratio (SNR)>4 (where SNR= ΔF/σ, and σ is the standard deviation of F_0_) were pooled to create an ensemble single-coverslip average. Coverslips with washout responses < 60% of control, indicating deteriorating cellular condition, were excluded from analysis. Values are shown as mean ± standard deviation (SD). ANOVA with Tukey’s *post hoc* test, Student unpaired t test, and 95% confidence intervals on the differences between mean values were used to determine statistical significance, defined as P<0.05. Data were tested for normality with the Shapiro-Wilk test.

## Results

### Electrical stimulation increases presynaptic cGMP

We characterized presynaptic cGMP dynamics in response to electrical stimulation using livecell imaging of the cGMP biosensor green cGull. Green cGull is a chimeric protein of the fluorescent protein citrine and mouse phosphodiesterase 5α that inceases fluorescence in response to micromolar cGMP with a 1000-fold selectivity over cyclic AMP^16^. Cultured rat hippocampal neurons were co-transfected with cGull and VAMP-mCherry, a chimera of red fluorescent protein and VAMP (vesicle associated membrane protein) as a marker of presynaptic boutons. **Figure 1A** shows basal and activity-driven cGull fluorescence compared to VAMP-mCherry fluorescence. Activity-dependent changes in cGMP levels were identified by electrical stimulation, using buffer supplemented with the AMPA and NMDA receptor antagonists CNQX and APV to prevent indirect excitation and isolate direct single neuron responses. Fluorescence increased in all cellular compartments in response to electrical stimulation, and was evident in presynaptic boutons (**Figure 1B)**.

**Figure 1.**
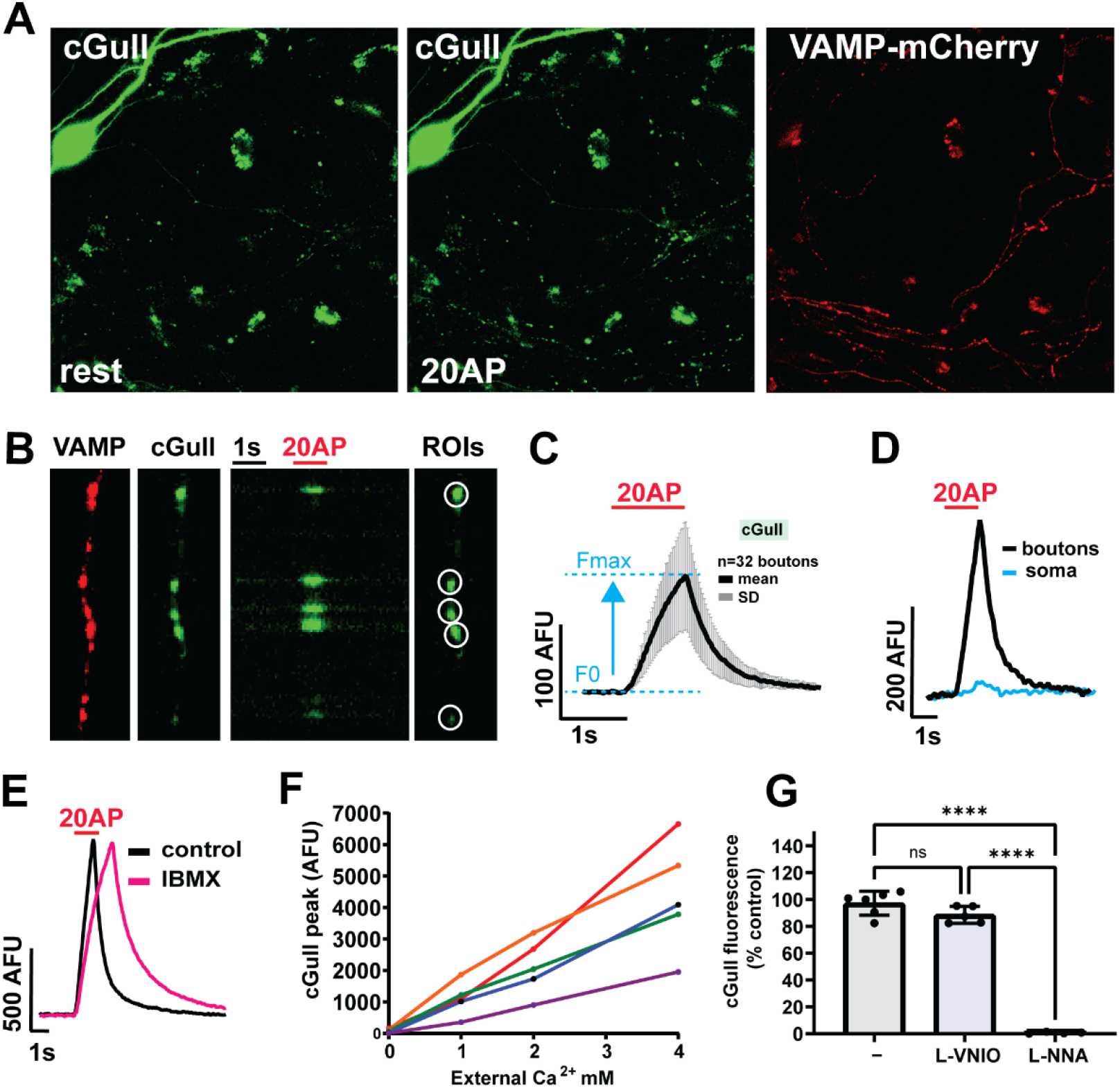
cGull reports basal and activity-driven cGMP concentrations. **(A**) At rest, cGull fluorescence is evident in somata and proximal dendrites (rest, left), and increases in all compartments following electrical stimulation (20 AP, middle), especially within presynaptic boutons co-expressing the presynaptic marker VAMP-mCherry (right). (**B**) Presynaptic boutons expressing VAMP-mCherry (VAMP) and cGull (far left and left), kymograph of evoked cGull fluorescence changes (middle), and ROIs drawn over responsive boutons (right). (**C**) Boutons from one cell analyzed together as an ensemble averaged time series. Peak amplitude: ΔF= Fmax-F_0_. Traces show mean over time, per cell. (**D)** Representative cGMP transients from boutons measured using cGull compared to signal from soma. (**E**) Transient waveform is prolonged in the presence of the phosphodiesterase inhibitor IBMX. (**F)** Presynaptic cGull transient peak amplitude increases with external Ca^2+^ concentration. **(G)** Presynaptic cGMP transients are resistant to the selective nNOS inhibitor L-VNIO, and completely abolished by the eNOS/nNOS inhibitor L-NNA. Calculated as percent of a control stimulation: control= 97.2± 8.9, n=6; L-VNIO = 88.6 ± 6.3, n= 5, L-NNA = 0.64 ± 0.60, n=4. Oneway ANOVA F(2, 12)= 272.3, p<0.0001 with Tukey’s *post hoc* test: control *vs* L-VNIO p=0.135, 95%CI [−2.41, 19.65], control *vs* L-NNA p<0.0001, 95%CI [84.8,108.3]; L-VNIO *vs* L-NNA p<0.0001, 95%CI [75.8, 100.2].

The time course of presynaptic cGMP dynamics indicated by cGull showed a rapid increase with a rapid decay (**Figure 1C)**. This presynaptic time course is consistent with the rapid sub-second kinetics of activity-driven NO/cGMP signaling reported in acutely isolated rat cerebellar cells^22^. Somatic cGMP did not change as markedly during electrical stimulation compared to presynaptic cGMP (**Figure 1D)**. Presynaptic cGMP transients showed prolonged signal decay in the presence of the phosphodiesterase-5 inhibitor IBMX, consistent with reduced cGMP hydrolysis (**Figure 1E)**.

NOS is activated by Ca^2+^, and many forms of NOS-mediated synaptic plasticity are dependent on Ca^2+^ influx through NMDA-type glutamate receptors ^1^. We also tested for NMDA receptor-independent presynaptic cGMP generation, which has been implicated in the regulations of HCN1 currents but not measured directly. Presynaptic cGull transient peak amplitude scaled with extracellular Ca^2+^ concentration in the presence of an NMDAR inhibitor (**Figure 1F**), suggesting that NMDAR-independent source of Ca^2+^ influx activates NOS, NO synthesis, and sGC.

We also tested the contributions of endothelial and neuronal NOS (eNOS and nNOS) to presynaptic cGMP transients. The cGMP transients detected by cGull were insensitive to the nNOS selective inhibitor L-VNIO but completely suppressed by the dual eNOS and nNOS inhibitor L-NNA (**Figure 1G**). Serial application of L-VNIO and L-NNA to the same cell showed that L-VNIO-resistant cGMP transients were blocked by L-NNA (**Supplemental Figure 1**).

### Volatile anesthetics depress cGMP transients

Live-cell imaging was used to test the effects of volatile anesthetics on electrically-evoked cGull transients following a pulse of 20 stimuli at 20 Hz (**Figure 2A**). CNQX and AP5 were included in all recording buffers to isolate single cell presynaptic changes in the absence recurrent excitation. Representative traces of presynaptic cGull transients in the presence of isoflurane or sevoflurane (0.51 ± 0.09 mM or 0.57± 0.13 mM, equivalent to 1.5 and 1.2 MAC, respectively) are shown in **Figure 2B**. Both isoflurane and sevoflurane inhibited presynaptic cGMP increases (**Figure 2C**). Peak cGMP transient amplitude was depressed by isoflurane (−33.2%; p<0.0001) or sevoflurane (−26.4%; p<0.0001) compared to control stimulation without anesthetic.

**Figure 2:**
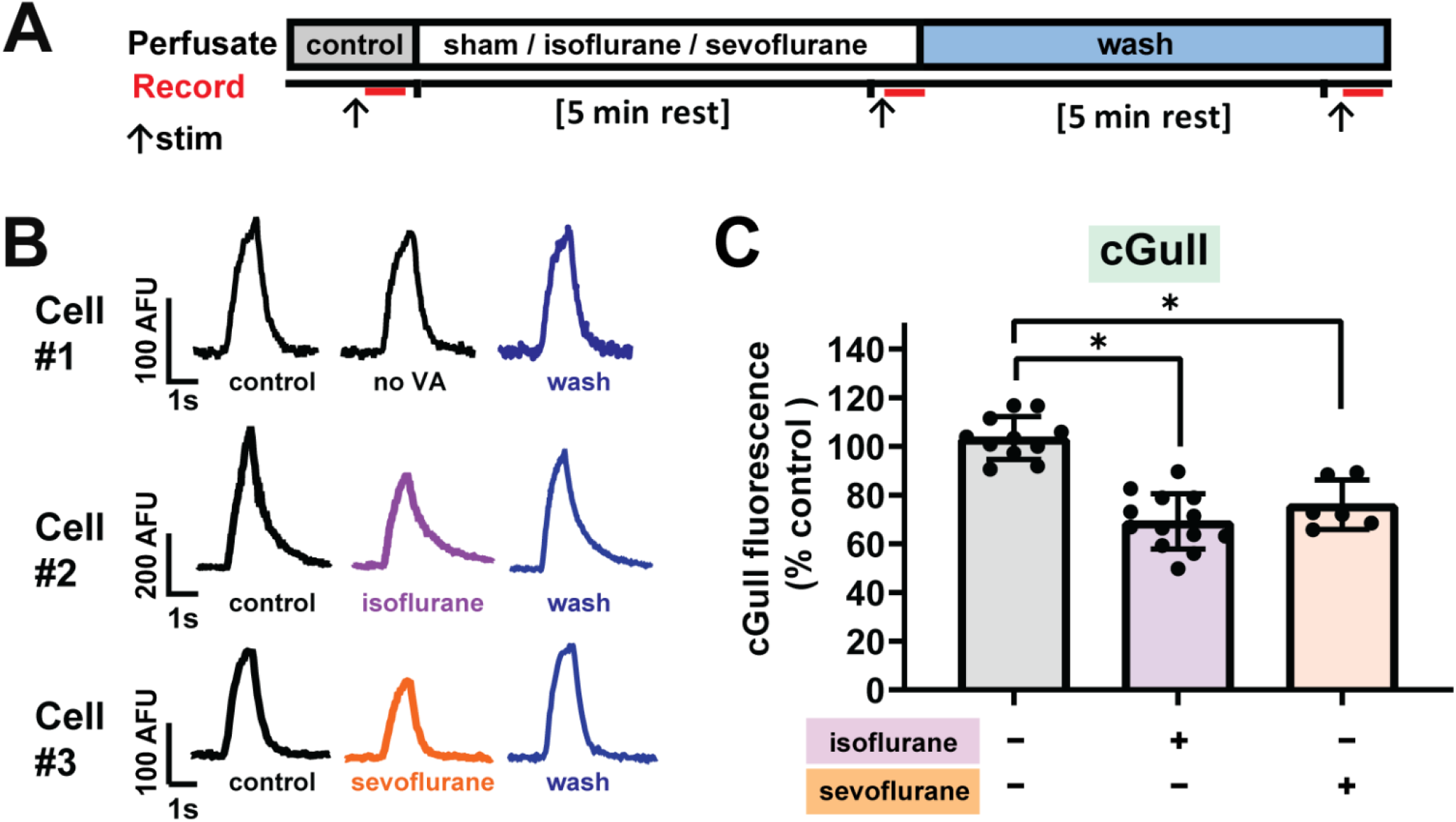
Volatile anesthetics inhibit stimulus-evoked presynaptic cGMP increases. **(A)** Live-cell imaging paradigm for electrical stimulation and drug perfusion paradigm. **(B**) Representative traces of presynaptic cGull transients in the presence of isoflurane or sevoflurane (0.51 ± 0.09 mM and 0.57 ± 0.13 mM). **(C)** Isoflurane or sevoflurane inhibits presynaptic cGMP increase, calculated as percent of control stimulation. Peak cGMP transient amplitude was depressed by both anesthetics compared to stimulation without anesthetic. Calculated as percent of control stimulation: no VA=103.6 ± 8.8 %, n=11; isoflurane=69.2 ± 11.3 %, n=13, sevoflurane = 76.2 ± 10.2 %, n=6. One-way ANOVA F(2, 27) = 35.3, p<0.0001, with Tukey’s *post hoc* test: no VA *vs* isoflurane p<0.0001, 95%CI [23.9, 44.7], no VA *vs* sevoflurane p<0.0001, 95%CI [14.5, 40.3]; isoflurane *vs* sevoflurane p=0.373, 95%CI [−19.4, −5.62].

### cGMP sequestration reduces isoflurane inhibition of presynaptic function

Since several cGMP effectors promote glutamatergic neurotransmission^1^, we hypothesized that reduced cGMP signaling by volatile anesthetics leads to impaired exocytosis. We analyzed synaptic vesicle (SV) exocytosis and presynaptic Ca^2+^ as measures of presynaptic function and excitability that are readily accessible to live-cell imaging. We used the genetically encoded cGMP scavenger protein sponGee in combination with optical biosensors for SV exocytosis (syn-pH) and presynaptic Ca^2+^ (syn-GCaMP6f). The recently developed sponGee is a chimeric protein of bovine cGMP-dependent protein kinase 1α and 1β binding sites, which has high affinity for cGMP but lack domains for downstream effector activation^17^. When cGull was cotransfected with sponGee (**Figure 3A)**, evoked presynaptic cGMP transients were completely blocked in all neurons observed, demonstrating that sponGee can scavenge electrically-evoked increases in presynaptic cGMP (**Figure 3B-C**).

**Figure 3:**
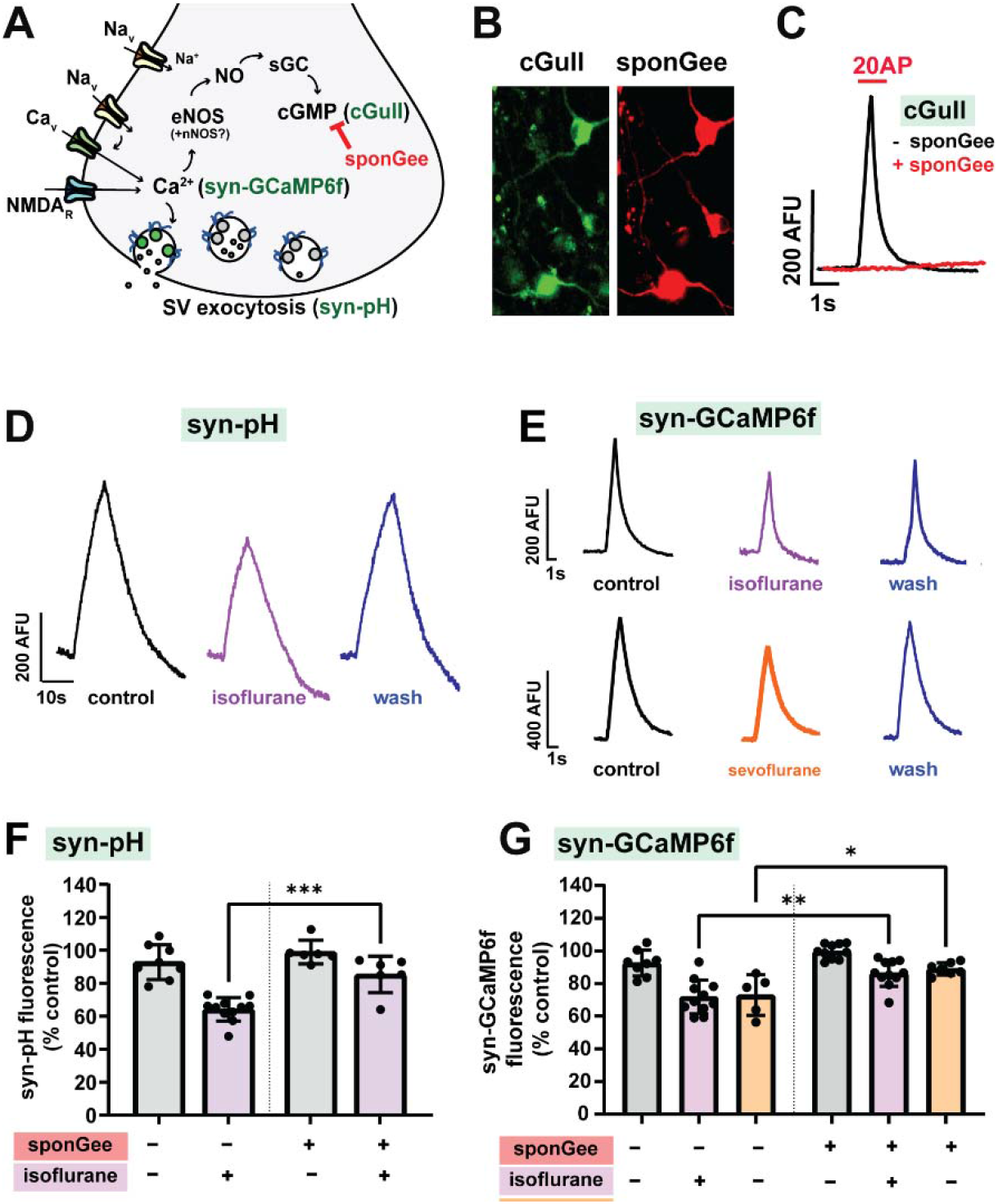
Sequestration of cGMP blunts volatile anesthetic inhibition of presynaptic function. **(A)** Probes used to investigate anesthetic effects on cGMP generation, Ca^2+^ influx, and synaptic vesicle (SV) exocytosis. Abbreviations: cGMP (cyclic guanosine monophosphate), eNOS and nNOS (endothelial and neuronal nitric oxide synthase), NO (nitric oxide), sGC (soluble gyanalyl cyclase), syn-GCaMP6f (synaptophysin-GCaMP6f), syn-pH (synaptophysin-pHluorin). **(B)** Overlapping red and green somatic fluorescence confirms sponGee-mRFP and cGull co-transfection. **(C)** Co-transfection of cGull with sponGee blocks activity-dependent cGMP increase. **(D)** Isoflurane reversibly inhibits SV exocytosis evoked by electrical stimulation as shown by representative traces from syn-pH transfected neurons, here −31.7%. **(E)** Isoflurane and sevoflurane reversibly inhibit presynaptic Ca^2+^ entry as shown by representative traces from syn-GCaMP6f transfected neurons, here −23.3 and −21.9% respectively. (**F**) Isoflurane inhibition of SV exocytosis is reduced by co-expression of syn-pH with the cGMP binding protein sponGee. Calculated as % control stimulation (left-right): syn-pH, 92.9 ± 10.6%, n=8; syn-pH plus isoflurane, 64.3 ± 7.1%, n=11; syn-pH and sponGee, 99.0 ± 7.2%, n=6; syn-pH and sponGee, plus isoflurane, 85.4 ± 11.1%. One-way ANOVA F(3,27)=25.84, p<0.0001, with Tukey’s *post hoc* test: syn-pH plus isoflurane *vs* syn-pH and sponGee plus isoflurane p=0.0004, 95%CI [8.71, 33.56]. (**G**) Isoflurane (and sevoflurane; data not shown) inhibition of presynaptic Ca^2+^ influx were both reduced by co-expressing syn-GCaMP6f with sponGee. Calculated as % control stimulation (left-right): syn-GCaMP6f = 92.4 ± 8.0%, n=8; syn-GCaMP6f plus isoflurane 71.8 ± 10.4%, n=11; syn-GCaMP6f plus sevoflurane 72.9 ± 12.4%, n=5; syn-GCaMP6f and sponGee 99.7 ± 4.5, n=10; syn-GCaMP6f and sponGee plus isoflurane 86.3 ± 8.0%, n=11; syn-GCaMP6f and sponGee plus sevoflurane 89.9 ± 3.0%, n=7. The following statistics are from one-way ANOVA F(5,46)=16.17, p<0.0001: Isoflurane: syn-GCaMP6f *vs* syn-GCaMP6f plus sponGee; p=0.0016, 95%CI [4.23, 24.83]. Sevoflurane: syn-GCaMP6f vs syn-GCaMP6f plus sponGee p=0.018, 95%CI [−30.18, −1.89].

We measured volatile anesthetic effects on presynaptic function using the paradigm shown in **Figure 2A**, with 20 AP at 20 Hz stimulation for syn-GCaMP6f and 100 AP at 10 Hz stimulation for syn-pH. Isoflurane reversibly inhibited presynaptic Ca^2+^ entry and synaptic vesicle exocytosis evoked by electrical stimulation (**3D-E**). Isoflurane inhibition of synaptic vesicle exocytosis was diminished by co-expression of the cGMP scavenger sponGee (p=0.0004, **Figure 3F**). Isoflurane inhibition of presynaptic Ca^2+^ influx was also diminished in neurons by coexpression of sponGee (p=0.0016, **Figure 3G**). This effect on Ca^2+^ influx was also seen with sevoflurane, another clinically used volatile anesthetic (p=0.018). Because sponGee affected both synaptic vesicle exocytosis and Ca^2+^ influx, we attribute the altered synaptic vesicle exocytosis to upstream effects on presynaptic excitability rather than direct cGMP actions on vesicular exocytosis mechanisms.

### Volatile anesthetic effect on presynaptic Ca^2+^ is PKG-independent

Multiple targets of cGMP can regulate presynaptic excitability. Cyclic nucleotide-gated (CNG) channels and hyperpolarization-activated cyclic nucleotide-gated (HCN) channels are directly modulated by cGMP, which increases channel activation^1^. Another set of channels depend on phosphorylation by cGMP-dependent protein kinase (PKG), including BK potassium channels and L-type voltage-gated Ca^2+^ channels. If PKG mediates cGMP effects on presynaptic excitability, a PKG inhibitor should recapitulate the effect of cGMP sequestration on presynaptic Ca^2+^ influx. We determined the effects of the PKG inhibitor KT-5823 on anesthetic inhibition of presynaptic Ca^2+^ influx (**Figure 4A).** KT-5823 had no effect on isoflurane inhibition of presynaptic Ca^2+^ (p=0.732, **Figure 4C**), which suggests that cGMP-dependent PKG targets do not contribute to isoflurane inhibition of activity-driven presynaptic Ca^2+^ influx.

**Figure 4:**
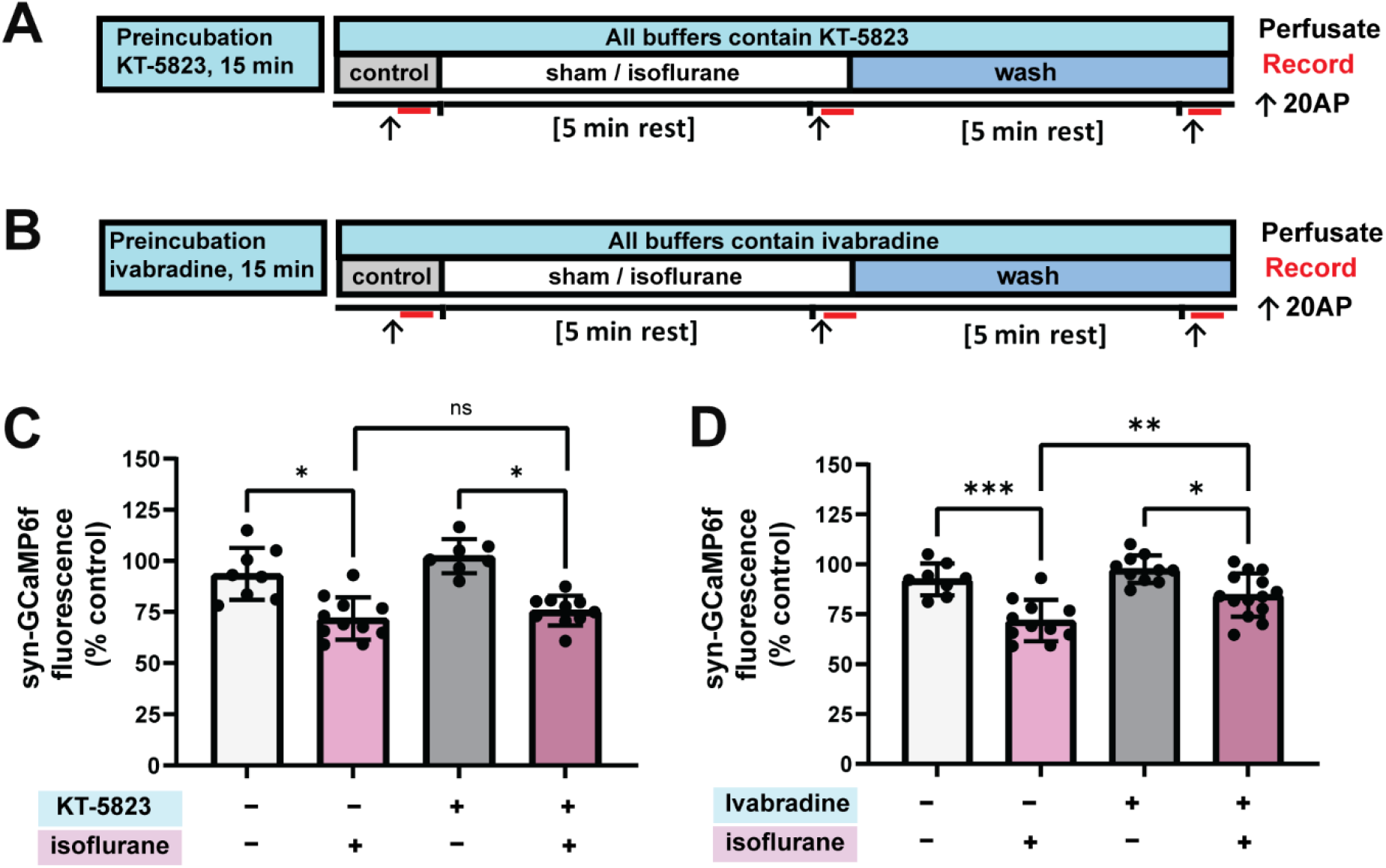
Isoflurane inhibits presynaptic Ca^2+^ influx via cGMP modulation of HCN channels independent of cGMP-dependent protein kinase. **(A)** Live-cell imaging paradigm to identify the contribution of cGMP-dependent protein kinase (PKG) to isoflurane actions, using the selective PKG inhibitor KT-5823. **(B)** Live-cell imaging paradigm to identify the contribution of HCN channels to isoflurane actions, using the HCN inhibitor ivabradine. **(C)** KT-5823 had no significant effect on isoflurane inhibition of presynaptic Ca^2+^ influx. Calculated as % control stimulation (left-right): no VA, 92.4 ± 8.0%, n=8; isoflurane, 71.8 ± 10.4%, n=11; KT-5823, no VA, 102.3 ± 8.4%, n=7; KT-5823 plus isoflurane, 75.7 ± 7.2%, n=10. One-way ANOVA, F(3,32)=18.58, with Tukey’s *post hoc* test: isoflurane *vs* isoflurane plus KT-5823, p=0.732, 95%CI [9.72, 31.61]**. (D)** Ivabradine diminished isoflurane inhibition of presynaptic Ca^2+^ influx. Calculated as % control stimulation (left-right): no VA, 92.4 ± 8.0%, n=8; isoflurane, 71.8 ± 10.4%, n=11; ivabradine, no VA, 97.6 ± 6.9%, n=10; ivabradine, isoflurane, 84.6 ± 10.8%, n=14. One-way ANOVA, F(3,38)=14.05, p<0.0001, with Tukey’s *post hoc* test: no VA vs isoflurane, p=0.00002, 95%CI [8.889, 32.43]; no VA, ivabradine *vs* isoflurane plus ivabradine, p=0.010, 95%CI [2.45, 23.4]; isoflurane *vs* isoflurane plus ivabradine, p=0.010, 95% CI [−23.0, −2.65].

### Volatile anesthetic effect on presynaptic Ca^2+^ influx is HCN-dependent

After excluding PKG-mediated mechanisms in the effects of cGMP on isoflurane actions, we analyzed the role of HCN channels, a known direct target of cGMP. HCN channels, major cGMP-modulated ion channels, contribute to presynaptic excitability and activity-dependent presynaptic Ca^2+^ influx by conducting a depolarizing mixed cation current. HCN channels are gated by cyclic nucleotides including cGMP, which shifts their voltage dependence of activation in the hyperpolarized direction to allow channel opening at more negative potentials. HCN/cGMP signaling is involved in depolarizing optic nerve axons ^23^ and increasing glutamate release probability in hippocampal neurons^21^. We hypothesized that volatile anesthetic depression cGMP synthesis and cGMP/HCN signaling would reduce HCN-mediated enhancement of presynaptic excitability. The general HCN inhibitor ivabradine diminished isoflurane inhibition of presynaptic Ca^2+^ influx (p=0.0087, **Figure 4B, D),** which supports a direct action of HCN in regulating presynaptic excitability and Ca^2+^ influx in a cGMP-dependent manner (**Supplemental Figure 2**).

## Discussion

We found that in rat hippocampal neurons, the volatile anesthetic isoflurane depresses presynaptic cGMP signaling to inhibit presynaptic Ca^2+^ influx and synaptic vesicle exocytosis. The mechanism involves inhibition of HCN channels, a downstream cGMP effector that normally sustains presynaptic excitability, through a reduction in simulation-evoked generation of cGMP.

### Presynaptic cGMP

We used the genetically encoded cGMP sensor cGull to characterize presynaptic cGMP dynamics during neurotransmission in cultured rat hippocampal neurons. Compared to biochemical assays, live-cell imaging of second messenger dynamics provides real-time compartment-specific resolution of the balance of cGMP synthesis and degradation with temporal and spatial resolution compared to previous static biochemical methods^24^.

We observed rapid presynaptic cGMP transients despite blockade of NMDA receptors, the best known mediator of Ca^2+^-dependent NO-sGC activation. External Ca^2+^ dependence of cGMP transient amplitude implicates another Ca^2+^ source supporting NO/sGC/cGMP signaling (i.e. voltage-gated Ca^2+^ channels). Our data suggest that NMDA receptor-independent NO synthesis depends on eNOS, and not nNOS, consistent with electrophysiological experiments showing that basal cGMP modulates glutamatergic signaling at a presynaptic site of action in an NMDA receptor-independent and eNOS-dependent manner^21^. As eNOS has been identified in the dendritic spines of primary cortical neurons^25^, one explanation is that dendritic NO generated by eNOS diffuses retrogradely to activate presynaptic sGC. In contrast, nNOS which is abundantly expressed in select interneurons, does not appear to participate, despite the presence of these interneurons in our primary cultures. The nNOS-mediated form of NO release is likely more NMDA receptor-dependent, as nNOS and NMDA receptor activation are spatially coupled by PSD95 tethering^26^.

### Anesthetic actions on presynaptic cGMP transients

The effects of isoflurane and sevoflurane were altered by sequestering cGMP and thereby inhibiting downstream cGMP signaling, achieved experimentally by co-expression of the cGMP-binding protein sponGee^17^. Previous investigations into anesthetic effects on NO/cGMP signaling show conflicting evidence of both inhibited and facilitated cGMP generation. This variability likely arises in part from differences in experimental paradigms to evoke neural activity and methods to quantify cGMP. Our observations are consistent with volatile anesthetic inhibition of cGMP in cultured cerebral neurons following NMDA application, even in the presence of the NMDA receptor blocker AP5^27^. In contrast, a recent study reported that sevoflurane increased cGMP in the forebrain of wild-type mice^7^. Because we measured presynaptic cGMP, for which there is no comparable study, further work is needed to resolve these apparently conflicting results, such as by live-cell imaging of somatic cGMP *in vivo*.

### Role of cGMP in depression of presynaptic excitability and neurotransmitter release

Previous studies have not identified downstream consequences of volatile anesthetic inhibition of cGMP synthesis on neuronal function. Two findings motivated us to investigate the role of cGMP in presynaptic anesthetic actions: 1) sensitivity to sevoflurane anesthesia is decreased in mice with a congenital deletion of sGC-1α, the isoform associated with presynaptic terminals,^7^ and 2) several cGMP effectors increase glutamate release^1, 28^.

Neurotransmitter release via synaptic vesicle (SV) exocytosis is steeply dependent on presynaptic Ca^2+^ influx through voltage-gated Ca^2+^ channels, itself determined by depolarization of the presynaptic terminal^29^. Isoflurane inhibition of presynaptic Ca^2+^ influx and transmitter release may involve action potential depression^30^ and diminished Ca^2+^ influx^31^. Previous work has identified presynaptic ion channel targets for volatile anesthetics including voltage-gated Na^+ 18^ and Ca^2+^ channels^32^, and “indirect” targets modulated by volatile anesthetic effects on second messenger signaling include α_2A_-adrenergic receptors^33^. Our current findings suggest that HCN channels may be an indirect target, modulated via volatile anesthetic effects on cGMP concentration.

### HCN channels as presynaptic effectors for anesthetic effects on cGMP

Our findings suggest that cGMP/HCN signaling partially mediates volatile anesthetic actions on presynaptic function in hippocampal neurons. Previous studies have implicated HCN in anesthetic mechanisms in thalamic^34^, thalamocortical^35^, and forebrain neurons^36^. HCN channels can be gated by either cAMP or cGMP; our study is the first to link volatile anesthetic depression of cGMP production with reduced HCN function. This effect may have been masked previously by direct volatile anesthetic actions on channel function^37^. Sequestration of cGMP supports a cGMP-dependent mechanism, however our studies do not exclude parallel cAMP-signaling effects as a regulator of HCN during volatile anesthetic exposure.

Our results are consistent with previous findings that presynaptic sGC1-mediated increases in cGMP increases glutamate release involving HCN channel activation in hippocampal glutamatergic neurons to increase presynaptic Ca^2+^ influx. ^21^ Our evidence for a role of HCN channels is indirect, so direct measurement of HCN currents is necessary to link reduced cGMP production to reduced channel function. Additional studies are also necessary to identify the specific HCN channel subtype(s) involved^38, 39^.

Within the hippocampus, HCN channel expression is best characterized in pyramidal neuron dendrites along a gradient with greatest enrichment in distal processes^40^. Axonal expression is not as well characterized, although HCN1 has been identified in axonal and presynaptic compartments of entorhinal neurons^41, 42^. Our experiments do not have the resolution to localize HCN1 or any particular HCN subtype in these cells to the active zone, but do indicate their functional role in regulating presynaptic excitability.

### Limitations and future experiments

A primary caveat to our proposed mechanism linking volatile anesthetic reductions in presynaptic cGMP levels, reduced HCN-mediated Ca^2+^ influx and synaptic vesicle exocytosis, and depressed presynaptic excitability is the absence of direct measurements of HCN channel function. Functional analysis is limited because cultures of dissociated neurons may not recapitulate the cellular properties and connectivity observed *in vivo*. Similarly, our focus on presynaptic function may exclude other mechanisms acting in an intact neuronal circuit, including glial contributions. We focused on presynaptic HCN channels because of their connections to presynaptic cGMP and Ca^2+^ dynamics related to exocytosis. However, this excludes the dendritic HCN pools, which likely contribute to cellular excitability *in vivo*^40, 43^. Further delineating the relationship between NO/CGMP signaling and anesthetic actions is challenging because of the complexity of overlapping signaling pathways within neurons. Several promising biosensors for NO may prove valuable in this regard ^**44, 45**^.

## Conclusions

We employed newly developed tools for studying volatile anesthetic effects on cGMP signaling that allow subcellular and single-cell resolution and manipulation of cGMP transients during neuronal activity and treatment. We observed that volatile anesthetics affect presynaptic cGMP/HCN signaling to inhibit Ca^2+^influx and neurotransmitter release. We provide the initial characterization of presynaptic cGMP dynamics during neuronal activity, which will be valuable in uncovering their relationships to presynaptic effects of anesthetics. We also observed cGMP transients generated independent of NMDA receptor activation involving eNOS. Building on this fundamental description of presynaptic cGMP dynamics, we observed a role for cGMP and HCN channels as mediators for volatile anesthetic modulation of presynaptic function.

## Acknowledgments

We thank members of the Hemmings laboratory for constructive comments.

---

**Supplementary Figure 1:**
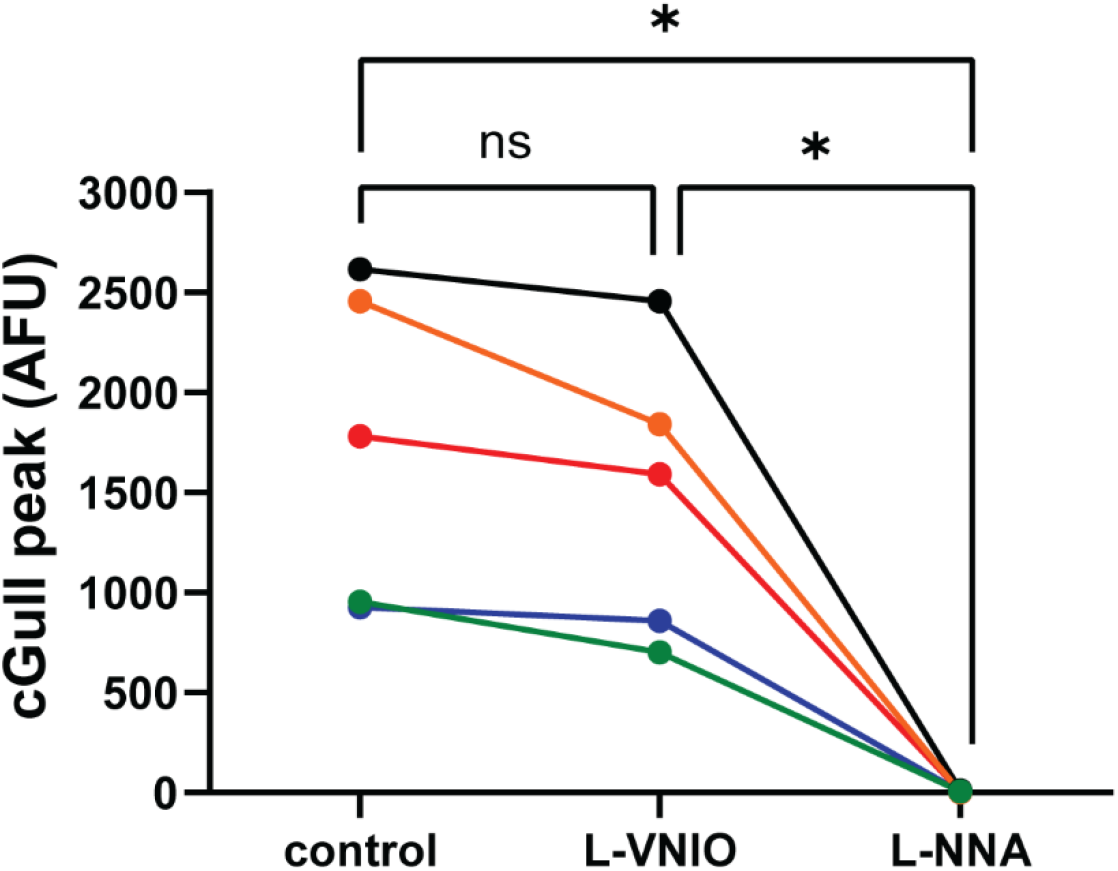
L-NNA depresses L-VNIO-resistant electrically-evoked presynaptic cGMP increase. Sequential perfusion of L-VNIO and L-NNA to the same cell was used to test the contribution of nNOS and eNOS to presynaptic cGMP transients (related to **Figure 1**). Distinct colors represent individual coverslips. One-way repeated measures ANOVA F(1.099, 4.395)=22.04, p=0.0070. Tukey’s *post hoc* test: control *vs* L-VNIO p=0.109, control *vs* L-NNA p=0.018, L-VNIO *vs* L-NNA p=0.022.

**Supplementary Figure 2:**
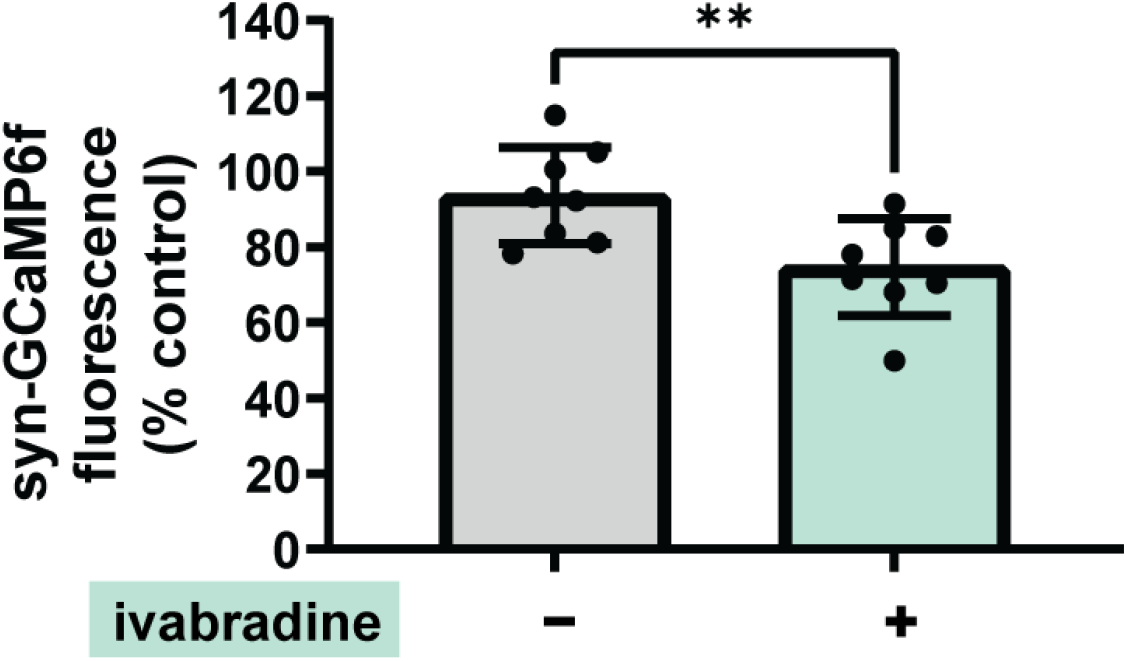
Ivabradine reduces presynaptic Ca^2+^ influx. The effect of ivabradine on presynaptic Ca^2+^ influx was measured in neurons expressing syn-GCaMP6f. Ca^2+^ transients evoked by 20 AP at 20 Hz stimulation were measured first without ivabradine, and then after a 20-min incubation with control buffer or with ivabradine. Ivabradine effect on evoked syn-GCaMP6f peak Ca^2+^ is displayed as percent of the control stimulation measured prior to drug incubation: Control= 93.6 ± 12.7%, n=8; ivabradine= 74.5 ± 12.8%, n=8; p=0.0100, 95%CI [−32.6, −5.3]. **, P < 0.05 by unpaired Student t-test.

## References

1 Hardingham N, Dachtler J, Fox K. The role of nitric oxide in pre-synaptic plasticity and homeostasis. Front Cell Neurosci 2013; 7:190

2 Toda N, Toda H, Hatano Y. Nitric oxide: involvement in the effects of anesthetic agents. Anesthesiology 2007; 107:822–42

3 Ichinose F, Huang PL, Zapol WM. Effects of targeted neuronal nitric oxide synthase gene disruption and nitroG-L-arginine methylester on the threshold for isoflurane anesthesia. Anesthesiology 1995; 83:101–8

4 Engelhardt T, Lowe PR, Galley HF, Webster NR. Inhibition of neuronal nitric oxide synthase reduces isoflurane MAC and motor activity even in nNOS knockout mice. Br J Anaesth 2006; 96:361–6

5 Johns RA, Moscicki JC, DiFazio CA. Nitric oxide synthase inhibitor dose-dependently and reversibly reduces the threshold for halothane anesthesia. A role for nitric oxide in mediating consciousness? Anesthesiology 1992; 77:779–84

6 Cechova S, Pajewski TN. The soluble guanylyl cyclase inhibitor ODQ, 1H-[1,2,4]oxadiazolo[4,3-a]quinoxalin-1-one, dose-dependently reduces the threshold for isoflurane anesthesia in rats. Anesth Anolg 2004; 99:752–7, table of contents

7 Nagasaka Y, Wepler M, Thoonen R, et al. Sensitivity to Sevoflurane anesthesia is decreased in mice with a congenital deletion of Guanylyl Cyclase-1 alpha. BMC Anesthesiol 2017; 17:76

8 Sjakste N, Baumane L, Meirena D, Lauberte L, Dzintare M, Kalvins I. Drastic increase in nitric oxide content in rat brain under halothane anesthesia revealed by EPR method. Biochem Pharmacol 1999; 58:1955–9

9 Berthiaume Y, Sapijaszko M, Mackenzie J, Walsh MP. Protein kinase C activation does not stimulate lung liquid clearance in anesthetized sheep. Am Rev Respir Dis 1991; 144:1085–90

10 Sjakste N, Sjakste J, Boucher JL, et al. Putative role of nitric oxide synthase isoforms in the changes of nitric oxide concentration in rat brain cortex and cerebellum following sevoflurane and isoflurane anaesthesia. Eur J Pharmacol 2005; 513:193–205

11 Nahrwold ML, Lust WD, Passonneau JV. Halothane-induced alterations of cyclic nucleotide concentrations in three regions of the mouse nervous system. Anesthesiology 1977; 47:423–7

12 Divakaran P, Rigor BM, Wiggins RC. Brain cyclic nucleotide responses to anesthesia with halothane delivered in air or purified oxygen. J Neurochem 1980; 35:514–6

13 Galley HF, Le Cras AE, Logan SD, Webster NR. Differential nitric oxide synthase activity, cofactor availability and cGMP accumulation in the central nervous system during anaesthesia. Br J Anaesth 2001; 86:388–94

14 Zuo Z, Tichotsky A, Johns RA. Inhibition of excitatory neurotransmitter-nitric oxide signaling pathway by inhalational anesthetics. Neuroscience 1999; 93:1167–72

15 Loeb AL, Gonzales JM, Reichard PS. Isoflurane enhances glutamatergic agonist-stimulated nitric oxide synthesis in cultured neurons. Brain Res 1996; 734:295–300

16 Matsuda S, Harada K, Ito M, et al. Generation of a cGMP Indicator with an Expanded Dynamic Range by Optimization of Amino Acid Linkers between a Fluorescent Protein and PDE5alpha. ACS Sens 2017; 2:46–51

17 Ros O, Zagar Y, Ribes S, et al. SponGee: A Genetic Tool for Subcellular and Cell-Specific cGMP Manipulation. Cell Rep 2019; 27:4003–12 e6

18 Speigel IA, Hemmings HC, Jr. Selective inhibition of gamma aminobutyric acid release from mouse hippocampal interneurone subtypes by the volatile anaesthetic isoflurane. Br J Anaesth 2021

19 Hoppa MB, Lana B, Margas W, Dolphin AC, Ryan TA. alpha2delta expression sets presynaptic calcium channel abundance and release probability. Nature 2012; 486:122–5

20 Solt K, Eger EI, 2nd, Raines DE. Differential modulation of human N-methyl-D-aspartate receptors by structurally diverse general anesthetics. Anesth Analg 2006; 102:1407–11

21 Neitz A, Mergia E, Eysel UT, Koesling D, Mittmann T. Presynaptic nitric oxide/cGMP facilitates glutamate release via hyperpolarization-activated cyclic nucleotide-gated channels in the hippocampus. Eur J Neurosci 2011; 33:1611–21

22 Bellamy TC, Garthwaite J. Sub-second kinetics of the nitric oxide receptor, soluble guanylyl cyclase, in intact cerebellar cells. J Biol Chem 2001; 276:4287–92

23 Garthwaite G, Bartus K, Malcolm D, et al. Signaling from blood vessels to CNS axons through nitric oxide. J Neurosci 2006; 26:7730–40

24 Sprenger JU, Nikolaev VO. Biophysical techniques for detection of cAMP and cGMP in living cells. Int J Mol Sci 2013; 14:8025–46

25 Caviedes A, Varas-Godoy M, Lafourcade C, et al. Endothelial Nitric Oxide Synthase Is Present in Dendritic Spines of Neurons in Primary Cultures. Front Cell Neurosci 2017; 11:180

26 Sattler R, Xiong Z, Lu WY, Hafner M, MacDonald JF, Tymianski M. Specific coupling of NMDA receptor activation to nitric oxide neurotoxicity by PSD-95 protein. Science 1999; 284:1845–8

27 Rengasamy A, Pajewski TN, Johns RA. Inhalational anesthetic effects on rat cerebellar nitric oxide and cyclic guanosine monophosphate production. Anesthesiology 1997; 86:689–98

28 Wang Q, Mergia E, Koesling D, Mittmann T. Nitric oxide/cGMP signaling via guanylyl cyclase isoform 1 modulates glutamate and GABA release in somatosensory cortex of mice. Neuroscience 2017; 360:180–9

29 Dittman JS, Ryan TA. The control of release probability at nerve terminals. Nat Rev Neurosci 2019; 20:177–86

30 Ouyang W, Hemmings HC, Jr. Depression by isoflurane of the action potential and underlying voltage-gated ion currents in isolated rat neurohypophysial nerve terminals. J Pharmacol Exp Ther 2005; 312:801–8

31 Baumgart JP, Zhou ZY, Hara M, et al. Isoflurane inhibits synaptic vesicle exocytosis through reduced Ca2+influx, not Ca2+-exocytosis coupling. Proc Natl Acad Sci U S A 2015; 112:11959–64

32 Koyanagi Y, Torturo CL, Cook DC, Zhou Z, Hemmings HC, Jr. Role of specific presynaptic calcium channel subtypes in isoflurane inhibition of synaptic vesicle exocytosis in rat hippocampal neurones. Br J Anaesth 2019; 123:219–27

33 Hara M, Zhou ZY, Hemmings HC, Jr. alpha2-Adrenergic Receptor and Isoflurane Modulation of Presynaptic Ca2+ Influx and Exocytosis in Hippocampal Neurons. Anesthesiology 2016; 125:535–46

34 Chen X, Sirois JE, Lei Q, Talley EM, Lynch C, 3rd, Bayliss DA. HCN subunit-specific and cAMP-modulated effects of anesthetics on neuronal pacemaker currents. J Neurosci 2005; 25:5803–14

35 Schwerin S, Kopp C, Pircher E, et al. Attenuation of Native Hyperpolarization-Activated, Cyclic Nucleotide-Gated Channel Function by the Volatile Anesthetic Sevoflurane in Mouse Thalamocortical Relay Neurons. Front Cell Neurosci 2020; 14:606687

36 Zhou C, Liang P, Liu J, et al. HCN1 Channels Contribute to the Effects of Amnesia and Hypnosis but not Immobility of Volatile Anesthetics. Anesth Analg 2015; 121:661–6

37 Riegelhaupt PM, Tibbs GR, Goldstein PA. HCN and K2P Channels in Anesthetic Mechanisms Research. Methods Enzymol 2018; 602:391–416

38 Shah MM. Cortical HCN channels: function, trafficking and plasticity. J Physiol 2014; 592:2711–9

39 He C, Chen F, Li B, Hu Z. Neurophysiology of HCN channels: from cellular functions to multiple regulations. Prog Neurobiol 2014; 112:1–23

40 Lorincz A, Notomi T, Tamas G, Shigemoto R, Nusser Z. Polarized and compartment-dependent distribution of HCN1 in pyramidal cell dendrites. Nat Neurosci 2002; 5: 1185–93

41 Bender RA, Kirschstein T, Kretz O, et al. Localization of HCN1 channels to presynaptic compartments: novel plasticity that may contribute to hippocampal maturation. J Neurosci 2007; 27:4697–706

42 Huang Z, Li G, Aguado C, Lujan R, Shah MM. HCN1 channels reduce the rate of exocytosis from a subset of cortical synaptic terminals. Sci Rep 2017; 7:40257

43 Ying SW, Jia F, Abbas SY, Hofmann F, Ludwig A, Goldstein PA. Dendritic HCN2 channels constrain glutamate-driven excitability in reticular thalamic neurons. J Neurosci 2007; 27:8719–32

44 Eroglu E, Gottschalk B, Charoensin S, et al. Development of novel FP-based probes for live-cell imaging of nitric oxide dynamics. Nat Commun 2016; 7:10623

45 Eroglu E, Charoensin S, Bischof H, et al. Genetic biosensors for imaging nitric oxide in single cells. Free Radic Biol Med 2018; 128:50–8

